## Supplementary Data for "Sea lamprey enlightens the origin of the coupling of retinoic acid signaling to vertebrate hindbrain segmentation"

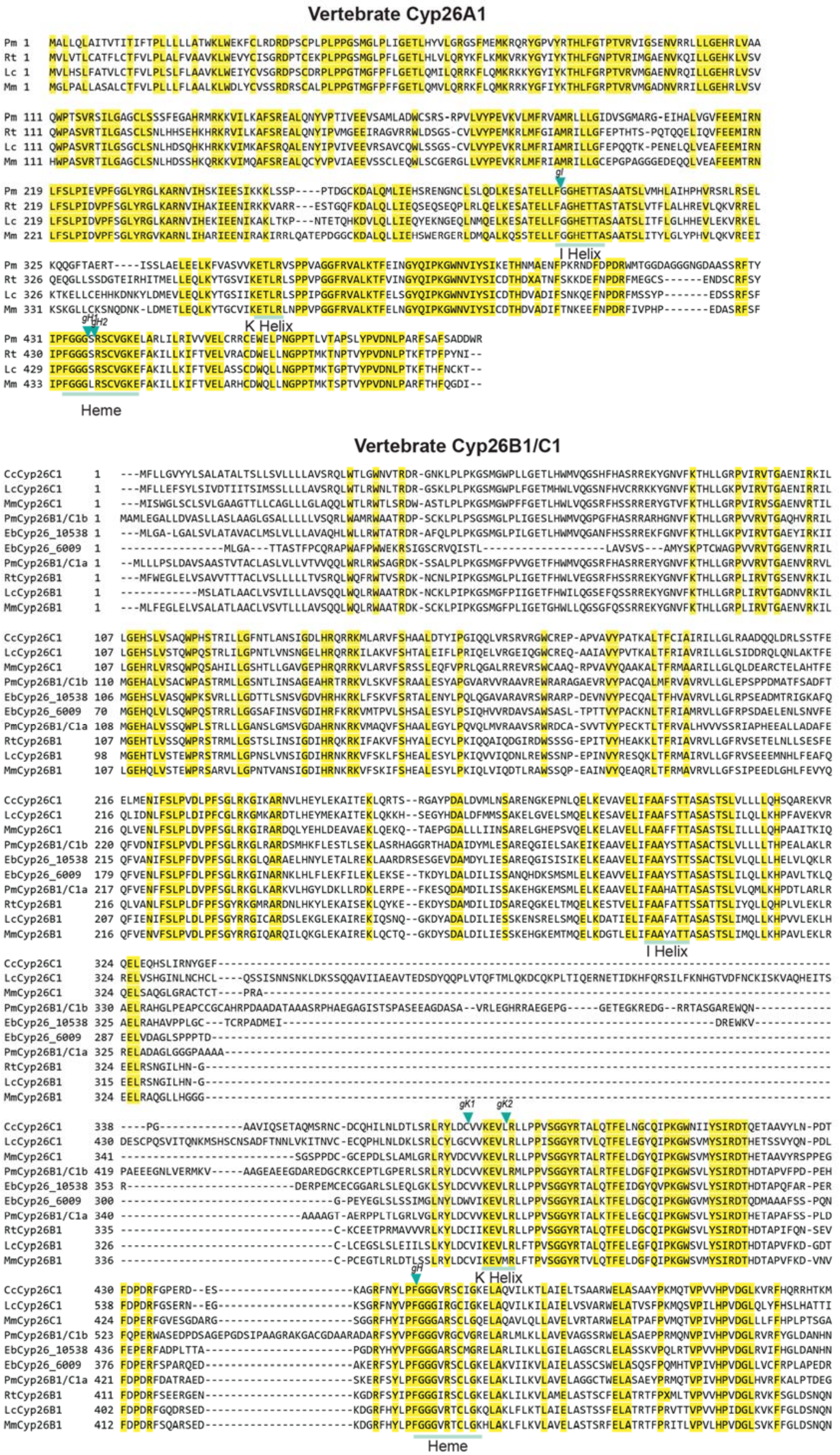
Vertebrate Cyp26A1
Pm
Rt
Lc
Mm
gI
I Helix
gH1
gH2
K Helix
Heme
Vertebrate Cyp26B1/C1
CcCyp26C1
LcCyp26C1
MmCyp26C1
PmCyp26B1/C1b
EbCyp26_10538
EbCyp26_6009
PmCyp26B1/C1a
RtCyp26B1
LcCyp26B1
MmCyp26B1
I Helix
gK1
gK2
gH
K Helix
Heme

**Supplementary Figure 1: Vertebrate Cyp26 protein alignments.** Protein alignments of vertebrate Cyp26A1 and vertebrate Cyp26B1/C1. The yellow highlighting represents the identity between sequences with a 95% threshold for sequence identity. Conserved K, I helices and Heme domains are indicated. Cutting sites of the gRNAs used for the CRISPR/Cas9 experiments are indicated with a teal arrow. Cc, Carcharodon carcharias; Eb, *Eptatretus burgeri*; Lc, *Latimeria chalumnae*; Mm, *Mus musculus*; Pm, *Petromyzon marinus*; Rt, *Rhincodon typus*.

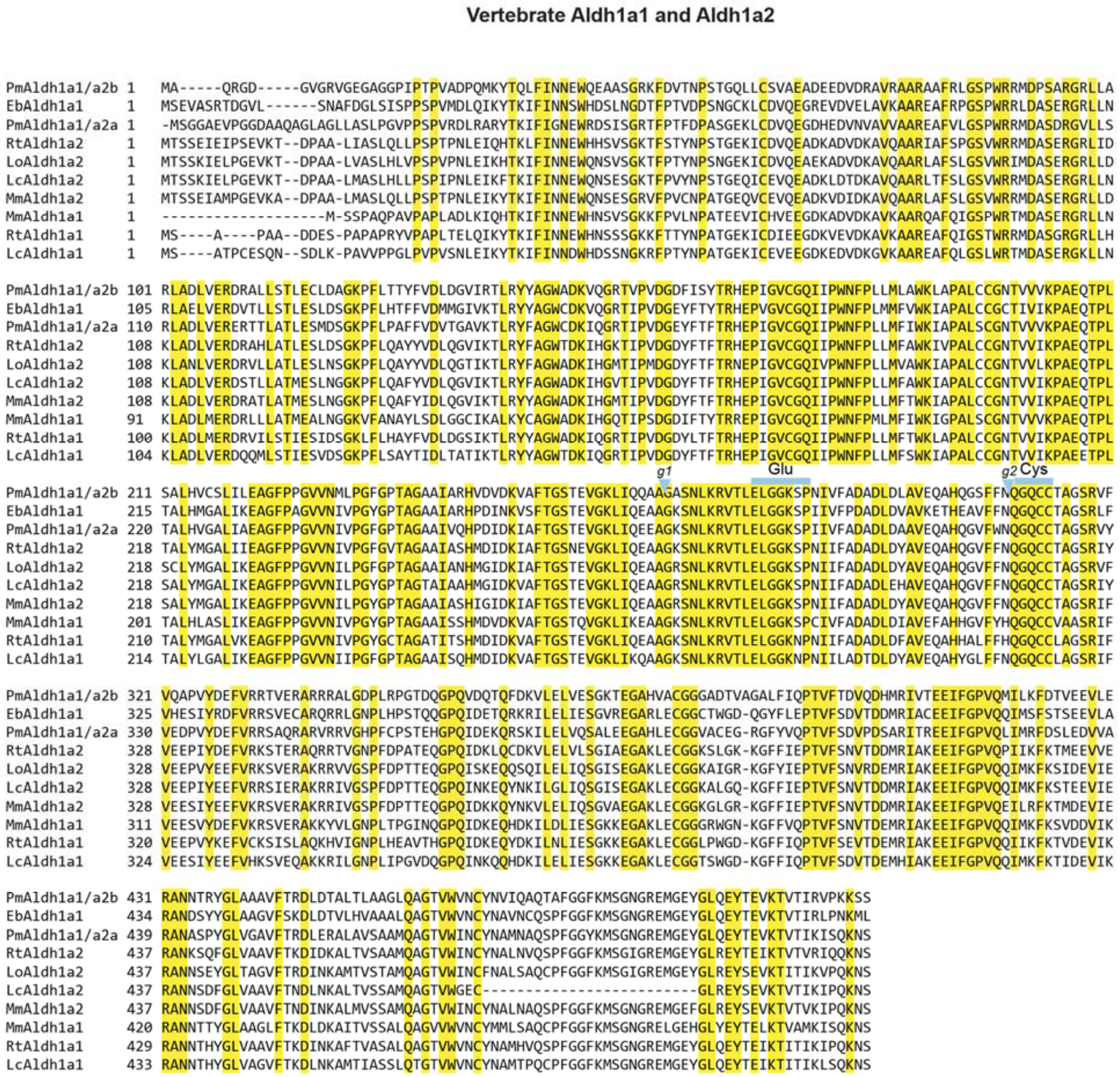

**Supplementary Figure 2: Vertebrate Aldh1a protein alignments.** Protein alignments of sea lamprey (Pm) Aldh1a1/a2a and Aldh1a1/a2b with Aldh1a1 and Aldh1a2 from various jawed vertebrates. The yellow highlighting represents the identity between sequences with a 95% identity threshold. Conserved Glutamic acid (Glu) and Cysteine (Cys) domains are indicated. Cutting sites of the gRNAs used for the CRISPR/Cas9 experiments are indicated with a blue arrow. Eb, *Eptatretus burgeri;* Lc, *Latimeria chalumnae*; Lo *Lepisosteus oculatus;* Mm, *Mus musculus;* Pm, *Petromyzon marinus*; Rt, *Rhincodon typus.*

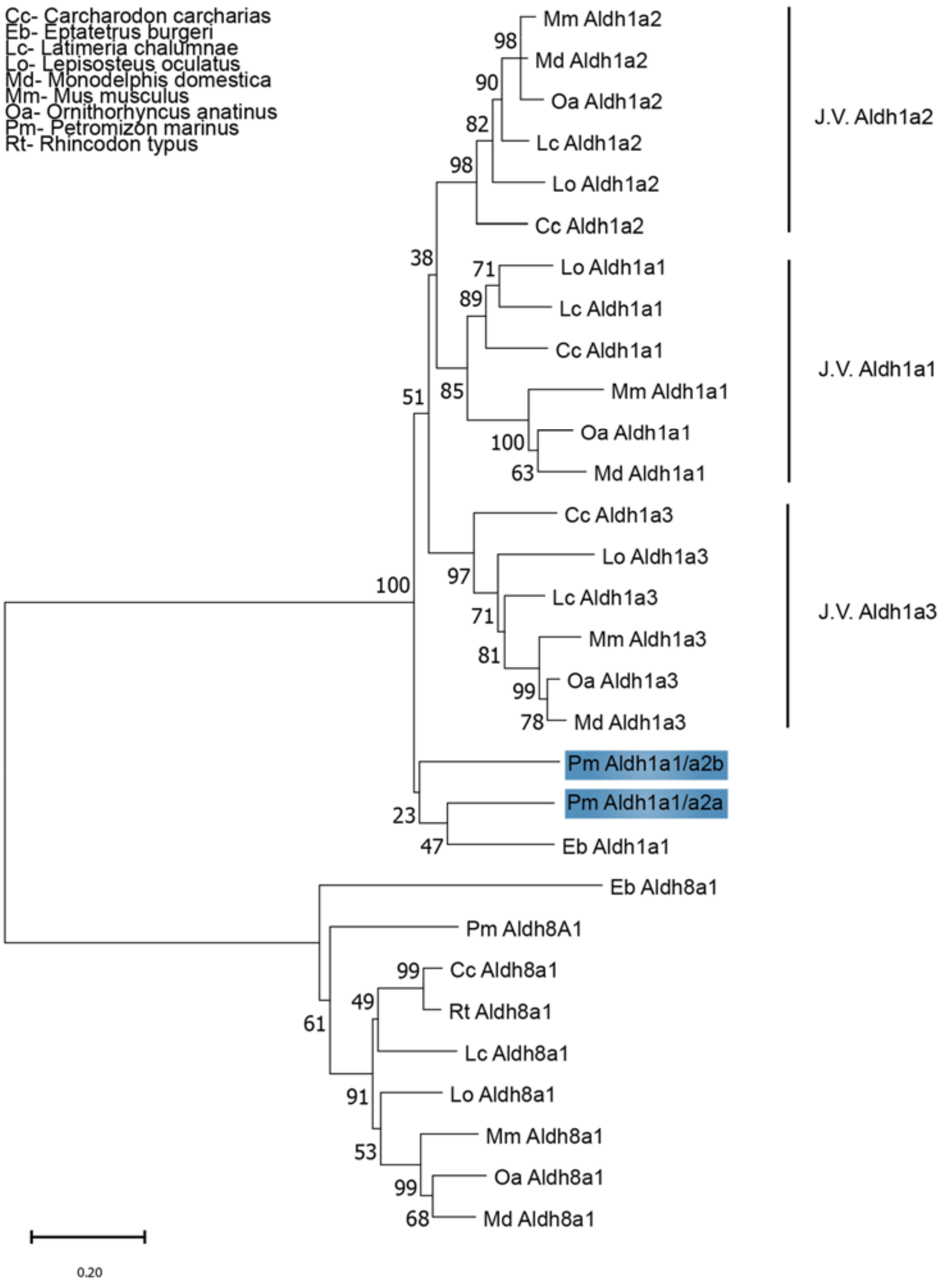

**Supplementary Figure 3: Vertebrate *Aldh1a* Phylogeny.** Phylogenetic analysis of the vertebrate *Aldh1a* complement with *Aldh1a8* used as an outgroup. Jawed Vertebrate (J.V.) *Aldh1a1/Aldh1a2/Aldh1a3* clades are indicated with black lines. Trees were generated by Maximum Likelihood using the WAG model with 500 iterations for bootstrap testing, and the resulting supporting value for each node is shown as a percentage. A scale bar for the evolutionary distance is indicated below each tree. Species name abbreviations are indicated on the top left of the panel.

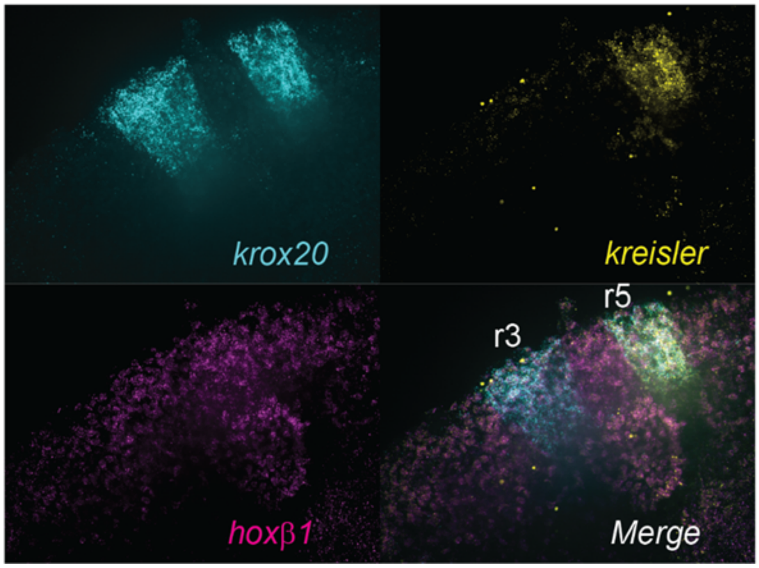

**Supplementary Figure 4: HCRv3 development and validation in the sea lamprey.** HCR panel representing a high-power (40X) image of *krox20* (cyan), *kreisler* (yellow) and *hoxβ1* (magenta) with their merged expression. This image was obtained after optimizing and adapting our HCRv3 protocol to be able to visualize hindbrain segments. We then used this protocol with this combination of genes to understand the changes in hindbrain segmentation in Talarazole vs. DMSO embryos (**Fig. 6a**).

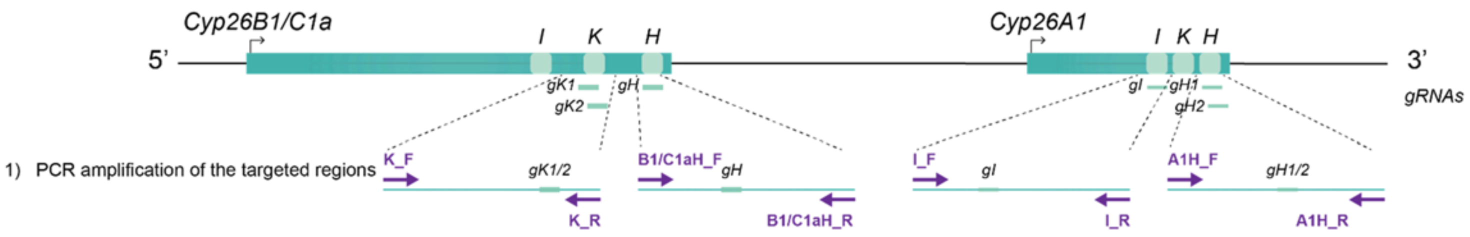

2) Sequencing of the targeted regions

```
Ctrl_C1H      GGAGCGTCCTCTACAGCATCCGCGACACGCACGAGACGGCGCCGGCCTTCAGCTCGCCGC   60
Cr_C1H_1      GGAGCGTCCTCTACAGCATCCGCGACACGCACGAGACGGCGCCGGCCTTCAGCTCGCCGC   60
Cr_C1H_2      GGAGCGTCCTGTACAGCATCCGCGACACGCACGAGACGGCGCCGGCCTTCAGCTCGCCGC   60
Cr_C1H_3      GGAGCGTCCTATACAGCATCCGCGACACGCACGAGACGGCGCCGGCCTTCAGCTCGCCGC   60
Cr_C1H_4      GGAGCGTCCTCTACAGCATCCGCGACACGCACGAGACGGCGCCGGCCTTCAGCTCGCCGC   60
Cr_C1H_5      GGAGCGTCCTCTACAGCATCCGCGACACGCACGAGACGGCGCCGGCCTTCAGCTCGCCGC   60
              ********** *************************************************

Ctrl_C1H      TTGACTTCGACCCCGACCGCTTCGACGCGACGCGCGCCGAGGACTCGAAGGAGCGCTTCA   120
Cr_C1H_1      TTGACTTCGACCCCGACCGCTTAGACGCGACGCGCGCCGAGGACTCGAAGGAGCGCTTCA   120
Cr_C1H_2      TTGACTTCGACCCCGACCGCTTCGACGCGACGCGCGCCGAGGACTCGAAGGAGCGCTTCA   120
Cr_C1H_3      TTGACTTCGACCCCGACCGCTTCGACGCGACGCGCGCCGAGGACTCGAAGGAGCGCTTCA   120
Cr_C1H_4      TTGACTTCGACCCCGACCGCTTCGACGCGACGCGCGCCGAGGACTCGAAGGAGCGCTTCA   120
Cr_C1H_5      TTGACTTCGACCCCGACCGCTTCG------------------------------------   69
              ************************

Ctrl_C1H      GCTACCTGCCGTTCGGCGGCGGCGTGCGCAGCTGCCTGGGCAAGGAGCTCGCCAAGCTCA   180
Cr_C1H_1      GCTACCTGCC----GGCGGCGGCGTGCGCAGCTGCCTGGGCAAGGAGCTCGCCAAGCTCA   176
Cr_C1H_2      GCTACCTGC----CGGCGGCGGCGTGCGCAGCTGCCTGGGCAAGGAGCTCGCCAAGCTCA   176
Cr_C1H_3      GCTACCTGC----CGGCGGCGGCGTGCGCAGCTGCCTGGGCAAGGAGCTCGCCAAGCTCA   176
Cr_C1H_4      GCTACCTGCC----GGCGGCGGCGTGCGCAGCTGCCTGGGCAAGGAGCTCGCCAAGCTCA   176
Cr_C1H_5      ---------------GCGGCGGCGTGCGCAGCTGCCTGGGCAAGGAGCTCGCCAAGCTCA   129
                             *********************************************

Ctrl_A1H      CCCGACCGCTGGATGACCGGCGGCGACGCCGGCGGCGGCAACGGAGACGCGGCATCGTCG   120
Cr_A1H_1      CCCGACCGCTGGATGACCGGCGGCGACGCCGGCGGCGGCAACGGAGACGCGGCATCGTC-   119
Cr_A1H_2      CCCGACCGCTGGATGACCGGCGGCGACGCCGGCGGCGGCAACGGAGACGCGGCATCGTCG   120
Cr_A1H_3      CCCGCCCGCGGGATGACCGGCGGCGCGGCCTGCGGCGGCATCGGAGACGTGGCATTGTCG   120
Cr_A1H_4      CCCGACCGCTGGATGACCGGCGGCGACGCCGGCGGCGGCAACGGAGACGCGGCATCGTC-   119
              **** **** ***************  *** ********* ******** ***** ***

Ctrl_A1H      CGCTTCACCTACATCCCGGGGGCGGCTCGCGCAGCTGCGTGGGCAAGGAGCTGGCGCGAC   180
Cr_A1H_1      ----------------------------GCGCAGCTGCGTGGGCAAGGAGCTGGCGCGAC   151
Cr_A1H_2      CGCTTCACCTCG--------------------CGCTGCGTGGGCAAGGAGCTGGCGCGAC   160
Cr_A1H_3      CGCGTCTC--------------------CTCCCGCTGCGTGGGCAAGGAGCTGGCGCGAC   160
Cr_A1H_4      ----------------------------GCGCAGCTGCGTGGGCAAGGAGCTGGCGCGAC   151
                                           ***************************

Ctrl_A1H      TCATCCTGAGGATCGTGGTGGTGGAGCTGTGCCGGCGCTGCGAGTGGGAGCTGCCCAACG   240
Cr_A1H_1      TCATCCTGAGGATCGTGGTGGTGGAGCTGTGCCGGCGCTGCGAGTGGGAGCTGCCCAACG   211
Cr_A1H_2      TCATCCTGAGGATCGTGGTGGTGGAGCTGTGCCGGCGCTGCGAGTGGGAGCTGCCCAACG   220
Cr_A1H_3      TCATCCTGAGGATCGTGGTGGTGGAGCTGTGCCGGCGCTGCGAGTGGGAGCTGCCCAACG   220
Cr_A1H_4      TCATCCTGAGGATCGTGGTGGTGGAGCTGTGCCGGCGCTGCGAGTGGGAGCTGCCCAACG   211
              ************************************************************

Ctrl_A1I      CCGCTCACCACTCCCGCCCCCTCCAGGACCTGAAGGAGTCGGCCACGGAGCTCCTGTTCG   420
Cr_A1I        CCGCTCACCACTCCCGCCCCCTCCAGGACCTGAAGGAGTCGGCCACGGAGCTCCTGT--G   419
              *********************************************************  *

Ctrl_A1I      GGGGACACGAGACGACGGCCAGCGCCGCCACGTCACTCGTCATGCACCTGGCAATCCACC   480
Cr_A1I        GGGGACACGAGACGACGGCCAGCGCCGCCACGTCGCTCGTCATGCACCTGGCAATCCACC   478
              ********************************** *************************
```

| | Cyp26A1_H | Cyp26A1_I | Cyp26B1/C1a_H |
|---|---|---|---|
| Mutant CRISPR loci | 4/7 | 1/12 | 5/6 |
| Control CRISPR loci | 5/5 | 6/6 | 3/3 |

**Supplementary Figure 5**: **Genotyping strategy and sequencing results for *Cyp26* CRISPR embryos**
Cartoon illustrating the design of primers for amplifying each site targeted by the *gRNAs* in the *Cyp26* loci. The primer binding sites are indicated in bold purple. For each site, we show an alignment of mutant loci (Cr_) sequences with the sequence of a control locus (Ctrl), the site targeted by the gRNA is indicated in bold and the PAM in red. A table indicating the proportion of mutant CRISPR or control CRISPR loci for each site is shown.

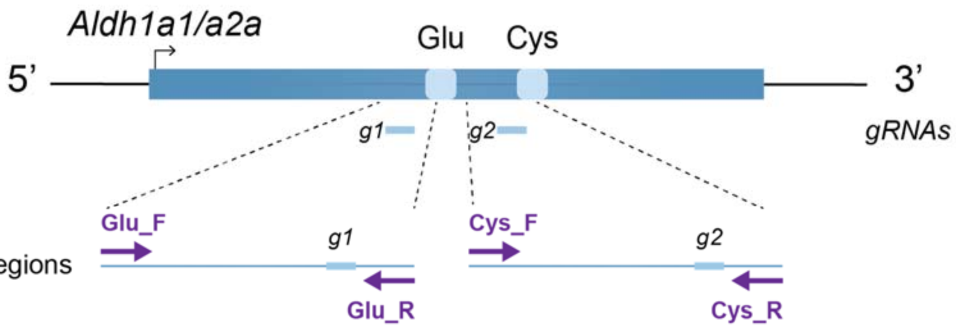

1) PCR amplification of the targeted regions

2) Sequencing of the targeted regions

```
Ctrl_Glu     CCCTTACGTAACGCAGGTGGGGAAGTTGATCCAGGAGGAGGCCGGCAAGAGTAACCTGAA   179
Cr_Glu_1     CCTTTACGTAACGCAGGTGGGGAAGTTGATCCAGGA--------------GTAACGTGAA   73
Cr_Glu_2     CCCTTACGTAACGCAGGTGGGGAAGTTGATCCAGGA--------------GTAACCTGAA   165
Cr_Glu_3     CCCTTACGTAACGCAGGTGGGGAAGTTGATCCAGGA--------------GTAACCTGAA   166
Cr_Glu_4     CCCTTACGTAACGCAGGTGGGGAAGTTGATCCAGGAGTAACCTGAAGC------------   165
Cr_Glu_5     CCCTTACGTAACGCAGGTGGGGAAGTTGATCCAGGA--------------GTAACCTGAA   165
             ** *********************************

Ctrl_Glu     GCGCGTGACGCTGGAGCTTGGCGGGAAGAGCCCCATCATCGTCTTCGCCGACGCCGACCG   239
Cr_Glu_1     GCGAGTGACGCTGAAGCTTGGCGGGAAGAGCCCCATCATCGTATTCGCCGAA-CCGACCG   132
Cr_Glu_2     GCGCGTGACGCTGGAGCTTGGCGGGAAGAGCCCCATCATCGTCTTCGCCGACGCCGACCG   225
Cr_Glu_3     GCGCGTGACGCTGGAGCTTGGCGGGAAGAGCCCCATCATCGTCTTCGCCGACGCCGACCG   226
Cr_Glu_4     ---GCGGACGCTGGAGCGTGGCGGGAAGAGCCCCATCATCGTTTTGACCGACGCCGACCG   222
Cr_Glu_5     GCGCGTGACGCTGGAGCTTGGCGGGAAGAGCCCCATCATCGTCTTCGCCGACGCCGACCG   225
                   ******* *** ************************ **  ****  *******

Ctrl_Cys     CCCCTGCTGTGACACTGCTGACTCGGCGTGGTTCCCACCCCTCCACCCCCCCCCTCCCAC   240
Cr_Cys_1     CCCCTGCTGTGACACTGCTGACTCGGCGTGGTTCCCACCCCTCCACCCCCCCCCTCCCAC   240
Cr_Cys_2     CCCCTGCTGTGACACTGCTGACTCGGCGTGGTTCCCACCCCTCCACCCCCCCCCTCCCAC   240
Cr_Cys_3     CCCCTGCTGTGACACTGCTGACTCGGCGTGGTTCCCACCCCTCCACCCCCCCC-TCCCAC   240
Cr_Cys_4     CCCCTGCTGTGACACTGCTGACTCGGCGTGGTTCCCACCCCTCCACCCCCCCCCTCCCAC   238
Cr_Cys_5     CCCCTGCTGTGACACTGCTGACTCGGCGTGGTTCCCACCCCTCCACCC------------   238
Cr_Cys_6     CCCCTGCTGTGACACTGCTGACTCGGCGTGGTTCCCACCCCTCCACCC------------   238
Cr_Cys_7     CCCCTGCTGTGACACTGCTGACTCGGCGTGGTTCCCACCCCTCCACCC------------   240
             ************************************************

Ctrl_Cys     TACGAGCAGTGGACGCGGCGGTGGAGCAGGCGCACCAGGGGGTGTTCTGGAACCAGGGCC   300
Cr_Cys_1     -------------------------------------------------------GGGCC   258
Cr_Cys_2     -------------------------------------------------------GGGCC   258
Cr_Cys_3     -------------------------------------------------------GGGCC   249
Cr_Cys_4     -------------------------------------------------------GGGCC   260
Cr_Cys_5     ------------------------------------------------------AGGGCC   250
Cr_Cys_6     ------------------------------------------------------AGGGCC   250
Cr_Cys_7     ------------------------------------------------------AGGGCC   244
                                                                    *****
```

| | Aldh1a1/a2a_Glu | Aldh1a1/a2a_Cys |
|---|---|---|
| Mutant CRISPR loci | 5/5 | 7/11 |
| Control CRISPR loci | 5/5 | 3/3 |

**Supplementary Figure 6**: **Genotyping strategy and sequencing results for *Aldh1a1/a2a* CRISPR embryos.** Cartoon illustrating the design of primers for amplifying each site targeted by the *gRNAs* in the *Aldh1a1/a2a* gene loci. Primer binding sites are indicated in bold purple. For each site, we show an alignment of mutant loci (Cr_) sequences with the sequence of a control locus (Ctrl), the site targeted by the gRNA is indicated in bold and the PAM in red. A table indicating the proportion of mutant CRISPR or control CRISPR loci for each site is shown.

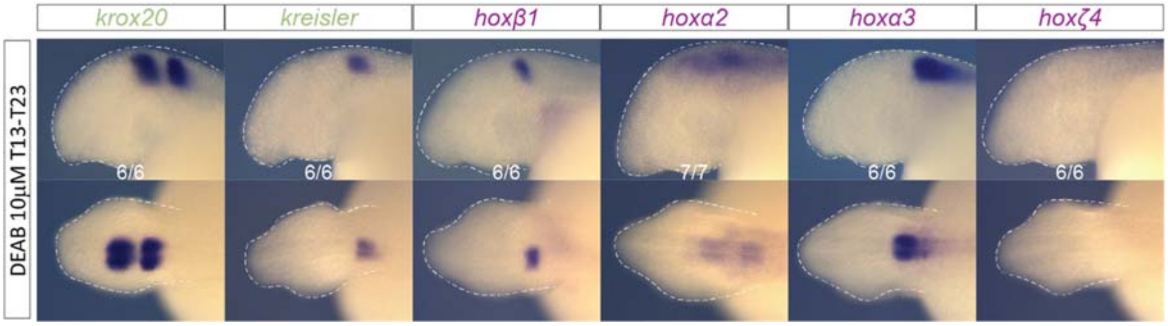

**Supplementary Figure 7**: **Effect of lower concentration of DEAB on hindbrain patterning.** cISH of key patterning hindbrain markers in embryos treated with 10 µM of DEAB at gastrulation (st13). For each gene, the most representative phenotype is shown, and numbers of experimental replicate are indicated. These molecular phenotypes are less severe than the ones obtained when treating the embryos with 50 µM DEAB (**Fig. 5d**) and are reminiscent of the phenotypes obtained in the Cr *Aldh1a1/a2a* mutants (**Fig. 8c**).

Mouse (*Mus musculus*): Aldh1a1: NP_001348432.1; Aldh1a2: NP_033048.2; Aldh8a1: NP_848828.1; Cyp26A1: XP_017173542.1; Cyp26B1: NP_001171184.1; Cyp26C1: XP_017173748.1; Cyp51: NP_064394.2

Grey short-tailed opossum (*Monodelphis domestica*): Aldh1a1: XP_001373128.2; Aldh1a2: XP_007479744.1; Aldh8a1: NP_848828.1; Cyp26A1: XP_001375292.2; Cyp26C1: XP_007478860.1; Cyp51-like: XP_001378190.2

Platypus (*Ornithorhynchus anatinus*): Aldh1a1: XP_007665209.1; Aldh1a2: XP_028925883.1; Aldh8a1: XP_001509667.1; Cyp26B1: XP_028902395.1; Cyp26C1: XP_028916075.1; Cyp51: XP_028926815.1

Spotted gar (*Lepisosteus oculatus*): Aldh1a1: XP_015222024.1; Aldh1a2: XP_006628784.1; Aldh8a1: XP_006625886.1; Cyp26A1-LG5: XP_015202021.1; Cyp26B1-LG5: XP_015202022.1; Cyp26B1-LG4: XP_006629157.1; Cyp51: XP_006636075.1

West indian coelacanth (*Latimeria chalumnae*): Aldh1a1: XP_006001725.2; Aldh1a2: XP_005998797.2; Aldh8a1: XP_014352052.1; Cyp26A1: XP_005991616.1; Cyp26B1: XP_005997719.1; Cyp26C1: XP_014341348.1; Cyp51: XP_014350652.1

Whale shark (*Rhincodon typus*): Aldh1a1-like: XP_015222024.1; Aldh1a2: XP_020369553.1; Aldh8a1: XP_048452951.1; Cyp26A1: XP_020382659.1; Cyp26B1: XP_020378695.1; Cyp26C1: XP_020367038.1; Cyp51: XP_020371022.1

Inshore hagfish (*Eptatretus burgeri*): Aldh1a1: ENSEBUG00000010361; Aldh8a1: ENSEBUG00000011145; Cyp26C1 (10538): ENSEBUG00000010538; Cyp26C1 (06909): ENSEBUG00000006909; Cyp51: ENSEBUG00000006843

Sea lamprey (*Petromyzon marinus*): Aldh1a1/a2a (Chr1): XP_032805864.1; Aldh1a1/a2b (Ch40): XP_032824682.1; Aldh8a1: XP_032810842.1; Cyp26A1: XP_032808019.1-2; Cyp26B1/C1a (Chr11): XP_032808017.1; Cyp26B1/C1b (Chr9): XP_032806163.1; Cyp51: XP_032816590.1

**Supplementary Figure 8: Protein accession numbers used for phylogeny analysis.** Protein sequences corresponding to each gene were retrieved using both ENSEMBL and NCBI databases, focusing on the latest available version of the genome and/or the most complete available genome annotation. Cyp51 and Aldh8a1 gene families were used as outgroups for the analyses of Cyp26 and Aldh1a complements, respectively.

Mouse (*Mus musculus*): *Aldh1a1*: NM_001361503.1; *Aldh1a2*: NM_009022.4; *Cyp26A1*: XM_017318053.2; *Cyp26B1*: NM_001177713.1; *Cyp26C1*: XM_017318259.3

Grey short-tailed opossum (*Monodelphis domestica*): *Aldh1a1*: XM_001373091.4; *Aldh1a2*: XM_007479682.2; *Cyp26A1*: XM_001375255.4; *Cyp26C1*: XM_007478798.1

Platypus (*Ornithorhynchus anatinus*): *Aldh1a1*: XM_007667019.3; *Aldh1a2*: XM_029070050.1; *Cyp26B1*: XM_029046562.2; *Cyp26C1*: XM_029060242.1

Chicken (*Gallus gallus*): *Aldh1a1*: NM_204577.5; *Aldh1a2*: NM_001397808.1; *Cyp26A1*: XM_015288598.4; *Cyp26B1*: XM_015286068.4; *Cyp26C1*: XM_421678.8

Spotted gar (*Lepisosteus oculatus*): *Aldh1a1*: XM_015366538.1; *Aldh1a2*: XM_006628721.2; *Cyp26A1LG5*: XM_015346535.1; *Cyp26B1-LG5*: XM_015346536.1; *Cyp26B1-LG4*: XM_006629094.2

West indian coelacanth (*Latimeria chalumnae*): *Aldh1a1*: XM_006001663.2; *Aldh1a2*: XM_005998735.2; *Cyp26A1*: XM_005991554.1; *Cyp26B1*: XM_005997657.2; *Cyp26C1*: XM_014485862.1

Whale shark (*Rhincodon typus*): *Aldh1a1-like*: XM_020527364.2; *Aldh1a2*: XM_020513964.2; *Cyp26A1*: XM_048608684.1; *Cyp26B1*: XM_020523106.1; *Cyp26C1*: XM_020511449.1

Sea lamprey (*Petromyzon marinus*): *Aldh1a1/a2a* (Chr1): XM_032949973.1; *Aldh1a1/a2b* (Chr40): XM_032968791.1; *Cyp26A1*: XM_032952128.1; *Cyp26B1/C1a* (Chr11): XM_032952126.1; *Cyp26B1/C1b* (Chr9): XM_032950272.1

**Supplementary Figure 9: mRNA accession numbers used for synteny analysis.** mRNA accession numbers of *Cyp26* and *Aldh1a* used to conduct the synteny analyses in different vertebrate models.

| Species | Exons | AA |
|---|---|---|
| *Pm Cyp26A1* | 7 exons | 497 AA |
| *Pm Cyp26B1/C1a* | 6 exons | 529 AA |
| *Pm Cyp26B1/C1b* | 6 exons | 634 AA |
| *Eb Cyp26 06909* | 6 exons | 468 AA |
| *Eb Cyp26 10538* | 6 exons | 528 AA |

**Supplementary Table 1: Cyp26 gene structure and protein length in cyclostomes.** Pm, *Petromyzon marinus* (sea lamprey); Eb, *Eptatretus burgeri* (inshore hagfish); AA. Amino-acid.

| Species | ***Cyp26A1*** | Cyp26A1 | ***Cyp26B1*** | Cyp26B1 | ***Cyp26C1*** | Cyp26C1 |
|---|---|---|---|---|---|---|
| Mm | 7 exons | 497 AA | 6 exons | 512 AA | 6 exons | 518 AA |
| Oa | NF | | 6 exons | 511 AA | 6 exons | 571 AA |
| Lo | 7 exons | 492 AA | 6 exons | 511 AA | 6 exons | 554 AA |
| Lc | 7 exons | 493 AA | 6 exons | 511 AA | 6 exons | 634 AA |
| Rt | 7 exons | 494 AA | 6 exons | 511 AA | NF | |
| Cc | | | | | 6exons | 560 AA |

**Supplementary Table 2: Cyp26 gene structure and protein length in jawed vertebrates.** Mm, *Mus musculus* (mouse); *Oa, Ornithorhynchus anatinus* (platypus)*; Lo, Lepisosteus oculatus* (spotted gar)*;* Lc, *Latimeria chalumnae* (coelacanth); Rt, *Rhincodon typus* (whale shark); *Cc, Carcharodon carcharias* (great white shark); NF: not found; AA. Amino-acid.

| Species | Exons | AA |
|---|---|---|
| *Pm Aldh1a1/a2a* | 13 | 520 AA |
| *Pm Aldh1a1/a2b* | 13 | 512AA or 459 AA |
| *Eb Aldh1a1* | 13 | 515 AA |

**Supplementary Table 3: Aldh1a gene structure and protein length in cyclostomes.** Pm, *Petromyzon marinus* (sea lamprey); Eb, *Eptatretus burgeri* (inshore hagfish); AA. Amino-acid.

| Species | ***Aldh1a1*** | Aldh1a1 | ***Aldh1a2*** | Aldh1a2 |
|---|---|---|---|---|
| Mm | 13 exons | 501 AA | 13 exons | 518 AA |
| Oa | 13 exons | 511 AA | 13 exons | 518 AA or 422 AA |
| Lo | 13 exons | 519 AA or 527 AA | 13 exons | 518 AA |
| Lc | 13 exons | 514 | 13 exons | 518 AA or 422 AA |
| Rt | 13 exons | 510 or 537 AA | 13 exons | 518 AA |

**Supplementary Table 4: Aldh1a gene structure and protein length in jawed vertebrates.** Mm, *Mus musculus* (mouse); *Oa, Ornithorhynchus anatinus* (platypus); *Lo, Lepisosteus oculatus* (spotted gar); Lc, *Latimeria chalumnae* (coelacanth); Rt, *Rhincodon typus* (whale shark); AA. Amino-acid.

| Gene | Accession Number | Probe set size | Amplifier/AlexaFluor | Average Probe Concentration |
|---|---|---|---|---|
| *Aldh1a1/a2a* | LOC116940317 | 19 | B3-488 | 6 nM |
| *Cyp26A1* | LOC116941264 | 23 | B4-546 | 5 nM |
| *Cyp26B1/C1a* | LOC116941263 | 22 | B1-647 | 5 nM |
| *krox20 (egr1)* | LOC116940673 | 18 | B3-488 | 6 nM |
| *kreisler (mafB-like)* | LOC116954108 | 12 | B4-546 | 11 nM |
| *hoxβ1* | LOC103091820 | 13 | B1-647 | 8 nM |

**Supplementary Table 5: HCR FISH probe set design.** Summary of the probe set design obtained for each gene targeted by FISH HCR; nM: nanomolar.

| name | 5'--> 3' sequence | description | product size |
|---|---|---|---|
| A1_H_F | TACAGCATCAAGGAGACGCA | Cyp26A1_H helix | 405 bp |
| A1_H_R | CGTCTTGTGCTGGCATAAATG | | |
| A1_I_F | GCTGAGACAAAGCGGGAAAG | Cyp26A1_I helix | 568 bp |
| A1_I_R | CCAGGCGAGGGAGATAACAT | | |
| **name** | **5'--> 3' sequence** | **description** | **product size** |
| C1_H_F | GGAGCGTCCTCTACAGCATC | Cyp26C1_H helix | 480 bp |
| C1_H_R | CTCCTCCGTCTCTCTGTCGT | | |
| **name** | **5'--> 3' sequence** | **description** | **product size** |
| Glu_F | TGGCGGTCGATTCTTCTCTT | Glutamate site genotyping | 360 bp |
| Glu_R | TATTGGCTGCGCAATCTCAC | | |
| Cys_F | ATATACAGCAGGCGTCAGGG | Cysteine site genotyping | 355 bp |
| Cys_R | CGCACGAACTCGTCGTAGA | | |

**Supplementary Table 6: Primers for CRISPR genotyping.** Primers used for genotyping targeted regions in CRISPR embryo; F: Forward, R: Reverse.
